# TBC1D18, a novel Rab5-GAP, coordinates endosome maturation together with Mon1

**DOI:** 10.1101/2021.11.11.468194

**Authors:** Shu Hiragi, Takahide Matsui, Yuriko Sakamaki, Mitsunori Fukuda

## Abstract

Endosome maturation is essential for efficient degradation of internalized extracellular molecules and plasma membrane proteins. Two Rab GTPases, Rab5 and Rab7, are known to regulate endosome maturation, and a Rab5-to-Rab7 conversion mediated by a Rab7 activator, Mon1–Ccz1, is essential for progression of the maturation process. However, the importance and mechanism of Rab5 inactivation during endosome maturation is poorly understood. Here we report a novel Rab5 inactivator (Rab5-GTPase activating protein [Rab5-GAP]), TBC1D18, which is associated with Mon1 and mediates endosome maturation. We found that Rab5 hyperactivation in addition to Rab7 inactivation occurs in the absence of Mon1. We present evidence showing that the severe defects in endosome maturation observed in *Mon1*-KO cells are attributable to Rab5 hyperactivation rather than to Rab7 inactivation. We then identified TBC1D18 as a Rab5-GAP by comprehensive screening of TBC-domain-containing Rab-GAPs. Expression of TBC1D18 in *Mon1*-KO cells rescued the defects in endosome maturation, whereas its depletion attenuated endosome formation and degradation of endocytosed cargos. Moreover, TBC1D18 was found to be able to interact with Mon1, and it localized in close proximity to lysosomes in a Mon1-dependent manner. Thus, TBC1D18 is a crucial regulator of endosome maturation that functions together with Mon1.

## INTRODUCTION

Endocytosis is a pivotal membrane dynamic process, in which cells internalize extracellular molecules and plasma membrane proteins (Sorkin & Zastrow, 2009; Sigismund et al., 2021). Invagination and fission of the plasma membrane results in the formation of early endosomes, where sorting of endocytosed cargos occurs. Some cargos are recycled back to the plasma membrane through recycling endosomes, whereas others are destined to lysosomes for degradation through the endocytic pathway in the following manner. Early endosomes first mature into late endosomes (also known as multivesicular bodies), a process that is accompanied by luminal acidification and acquisition of lysosomal enzymes from the *trans-Golgi* network, and the late endosomes then fuse with lysosomes. Endocytosed cargos are eventually degraded in the lysosomes, and the products of degradation are transported out of the lysosomes for energy generation or reutilization in biosynthetic pathways.

Rab GTPases are widely thought to coordinate protein sorting and vesicle/membrane trafficking during the endosome maturation processes. Rabs function as a molecular switch by cycling between an inactive form (GDP-bound) and active form (GTP-bound). Although inactive Rabs are present in the cytosol, active Rabs are localized to specific membrane compartments (or organelles) via their C-terminal cysteine residue(s) modified by geranylgeranyl transferase and recruit a specific effector protein(s) that promotes vesicle/membrane trafficking (Stenmark, 2009; Hutagalung & Novick, 2011; Pfeffer, 2017; Homma et al., 2021). Rab activity is thought to be spatiotemporally regulated by activation and inactivation enzymes, i.e., guanine nucleotide exchange factors (GEFs) and GTPase-activating proteins (GAPs), respectively (Lamber et al., 2019). Two Rab family members, early endosomal Rab5 (Ypt5 in yeasts) and late endosomal Rab7 (Ypt7 in yeasts) are key Rabs that coordinate endosome maturation (Huotari & Helenius, 2011; Borchers et al., 2021). After endocytosis, Rab5 is activated by a specific GEF, such as Rabex5 (Horiuchi et al., 1997), and it facilitates the formation of early endosomes and their homotypic fusion (Stenmark et al., 1994). The early endosomal Rab5 then directly recruits the Mon1–Ccz1 heterodimeric complex, a Rab7-GEF, to the same compartment (Nordmann et al., 2010; Kinchen & Ravichandran, 2011), and the recruited Mon1–Ccz1 activates Rab7, which regulates subsequent late endosome maturation. The Rab7 on late endosomes recruits the hexameric homotypic fusion and vacuole protein sorting (HOPS) tethering complex, which mediates fusion between late endosomes and lysosomes (Balderhaar & Ungermann, 2013). Finally, the late endosomal Rab7 is inactivated by a GAP, such as TBC1D15 (Zhang et al., 2005), and reused for the next round of endosome maturation.

The spatiotemporal regulation of Rab5 and Rab7 activities by their GEFs and GAPs is important, because the Rab5-to-Rab7 conversion described above is essential for endosome maturation. While involvement of a Rab5-GEF, Rab7-GEF, and Rab7-GAP in endosome maturation has been well studied, very little has been known about the importance of Rab5 inactivation or the involvement of a specific Rab5-GAP in the Rab5-to-Rab7 conversion. In this study, we identified a novel Rab5-GAP, TBC1D18, that mediates endosome maturation. We discovered that *Mon1* knockout (unless otherwise specified, double KO [DKO] of *Mon1a* and *Mon1b*, i.e., *Mon1a/b*-DKO is referred to simply as *Mon1*-KO below) in mammalian cells results in hyperactivation of Rab5, even though Rab7 is inactivated, suggesting that Mon1 is involved not only in Rab7 activation but also in Rab5 inactivation as well. We also showed that expression of TBC1D18 is capable of rescuing defects in endosome maturation in *Mon1*-KO cells, whereas its depletion results in enlargement of early endosomes and attenuated degradation of endocytosed cargos. Based on our findings, we propose a new endosome maturation model in which Mon1–Ccz1 together with TBC1D18 coordinates endosome maturation by inactivating Rab5 and activating Rab7.

## RESULTS

### Distinct phenotypes of *Mon1*-KO cells and *Rab7*-KO cells in endosome maturation

To investigate the function of Mon1 in endosome maturation in greater detail, we first established *Mon1*-KO (*Mon1a/b*-DKO) MDCK cells (Fig. S1A) and examined their lysosomal morphology by immunostaining for lysosomal membrane-associated protein 2 (LAMP2). Although, consistent with our previous observation, the *Rab7*-KO cells contained slightly larger lysosomes than WT cells did (Kuchitsu et al., 2018), the *Mon1*-KO cells contained extremely enlarged lysosomes (Fig. 1A) (van den Boomen et al., 2020). The *Mon1*-KO phenotype could not have been caused by an off-target effect of guide RNAs, because it was completely rescued by re-expression of either Mon1a or Mon1b (Fig. 1A), thereby suggesting that Mon1a and Mon1b function redundantly in endosome maturation. Actually, KO of *Mon1a* or *Mon1b* alone had no effect on lysosomal morphology (data not shown). Moreover, *Mon1*-KO resulted in significantly fewer LAMP2-positive dots in comparison with the WT and *Rab7*-KO cells (Fig. 1B and 1C). Similar observations, i.e., enlarged size and reduced numbers, were also observed in regard to early endosomes (early endosome antigen 1 [EEA1]) and late endosomes (lysobisphosphatidic acid [LBPA]) in the *Mon1*-KO cells, but Golgi morphology (GM130, a *cis*-Golgi marker) was unaltered (Fig. S1B and S1C). Since all of these *Mon1*-KO phenotypes were also observed in *Ccz1*-KO cells (data not shown), the Mon1–Ccz1 complex is required for the proper morphology of both endosomes and lysosomes (van den Boomen et al., 2020). Although the lysosomes in the *Mon1*-KO cells were considerably enlarged, they appeared to have acquired a highly acidic pH and active lysosomal proteases, because they stained positive with both LysoTracker and Magic Red, which label acidic organelles and lysosomal protease activity, respectively (Fig. S1D).

**Figure 1.**
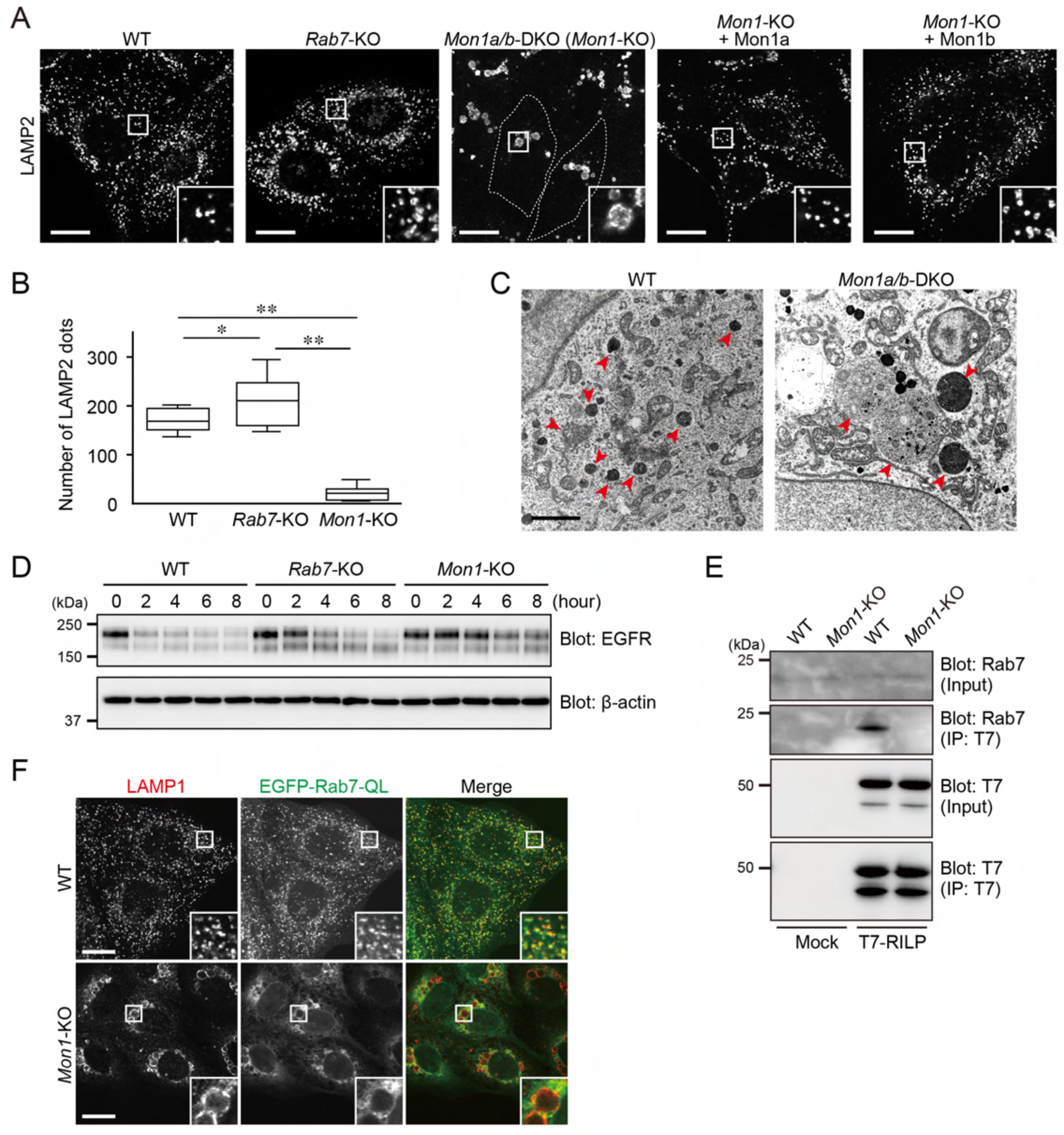
Severe defects in the endosome maturation of *Mon1*-KO cells. (A) Immunostaining of LAMP2 in the cells indicated. *Mon1*-KO cells are outlined with dotted lines. Insets show magnified views of the boxed areas. Scale bars, 20 μm. (B) The numbers of LAMP2-positive dots per cell (n = 15 cells) were counted. The solid bars, boxes, and whiskers indicate the median, interquartile range (25th to 75th percentile), and upper and lower quartiles, respectively, of the values. The statistical analyses were performed by one-way ANOVA and Tukey’s test. *, *P* <0.05; **, *P* <0.01. (C) WT and *Mon1*-KO cells were examined by electron microscopy. The red arrowheads point to MVBs and lysosomes. Scale bars, 1 μm. (D) Degradation of EGFR in the indicated cells after exposure to 200 ng/mL EGF. Their cell lysates were analyzed by immunoblotting with the antibodies indicated. (E) The amount of active, GTP-bound Rab7 in WT and *Mon1*-KO cells was analyzed by an effector-based pull-down assay with beads coupled to T7-RILP (Cantalupo et al., 2001). (F) Immunostaining of LAMP1 in WT and *Mon1*-KO cells stably expressing EGFP-Rab7-QL. Insets show magnified views of the boxed areas. Scale bars, 20 μm.

We next investigated whether substrates of endocytosis can be degraded in *Mon1*-KO cells. As shown in Fig. 1D, ligand (epidermal growth factor [EGF])-induced degradation of EGF-receptor (EGFR) was substantially attenuated in *Mon1*-KO cells in comparison with its degradation in WT and *Rab7*-KO cells. Similarly, fluorescence signals of DQ-bovine serum albumin (BSA), which is taken up by fluid-phase endocytosis, were clearly more reduced in *Mon1*-KO cells than in WT and *Rab7*-KO cells (Fig. S1E). It was noteworthy that the maturation of endosomes that contained substrates of endocytosis was more strongly inhibited by Mon1 depletion than by Rab7 depletion, suggesting that Mon1 has another function(s) in addition to Rab7 activation during endosome maturation. We therefore attempted to identify the new role, presumably the Rab7-independent role, of Mon1 in endosome maturation by analyzing the phenotypes of *Mon1*-KO cells in greater detail.

### Hyperactivation of Rab5, not inactivation of Rab7, in *Mon1*-KO cells induced formation of greatly enlarged endosomes and lysosomes

Because Mon1 is known to be a component of the Rab7-GEF complex (Kinchen & Ravichandran, 2010; Nordmann et al., 2010; Poteryaev et al., 2010), we first checked the activity of Rab7 in WT and *Mon1*-KO cells by performing effector pull-down assays with beads coupled to T7-tagged RILP, the best characterized Rab7 effector (Cantalupo et al., 2001). After incubating lysates of WT and *Mon1*-KO cells with the T7-RILP beads, the active Rab7 trapped by the beads was analyzed by immunoblotting (Fig. 1E). As anticipated, the level of active Rab7 in the *Mon1*-KO cells was markedly reduced in comparison with its level in the WT cells. To determine whether inactivation of Rab7 directly caused the greatly enlarged lysosomes and endosomes (lysosomes/endosomes) in *Mon1*-KO cells, we overexpressed a constitutively active Rab7 mutant with an EGFP-tag (EGFP-Rab7-QL) in *Mon1*-KO cells (Fig. 1F). However, the results showed that greatly enlarged lysosomes were observed in *Mon1*-KO cells even in the presence of an excess amount of active Rab7, suggesting that inactivation of Rab7 alone is unlikely to induce the phenotypes observed in *Mon1*-KO cells.

We next turned our attention to Rab5, which functions upstream of Mon1, because a previous study had reported that overexpression of a constitutively active Rab5A mutant, Rab5A-QL, in cultured mammalian cells induced greatly enlarged lysosomes/endosomes (Wegener et al., 2010) (Fig. S2A), a phenotype that resembled the phenotypes of the *Mon1*-KO cells that we had observed in this study. To determine whether the *Mon1*-KO phenotypes were caused by hyperactivation of Rab5, like Rab5-QL overexpression, we simultaneously knocked down endogenous Rab5A/B/C in WT and *Mon1*-KO cells with specific siRNAs and then immunostained them for EEA1 and LAMP2 (Figs. S2B and 2A). Upon depletion of all three Rab5 isoforms, highly enlarged endosomes/lysosomes were no longer observed in the *Mon1*-KO cells, and the number of LAMP2-positive dots was restored (Fig. 2B). Furthermore, depletion of Mon1 facilitated activation of Rab5A as revealed by active Rab5 pull-down assays with glutathione *S*-transferase (GST)-tagged Appl1-N (Zhu et al., 2007) (Fig. 2C). Consistent with these results, EGFP-tagged Rab5A was clearly associated with the limiting membrane of the enlarged lysosomes/endosomes in the *Mon1*-KO cells (Figs. 2D and S2C). Taken together, these results indicated that the phenotypes of *Mon1*-KO cells depend on hyperactivation of Rab5, not inactivation of Rab7, and implied that Mon1 has an inhibitory role in relation to Rab5, that is, inactivates Rab5.

**Figure 2.**
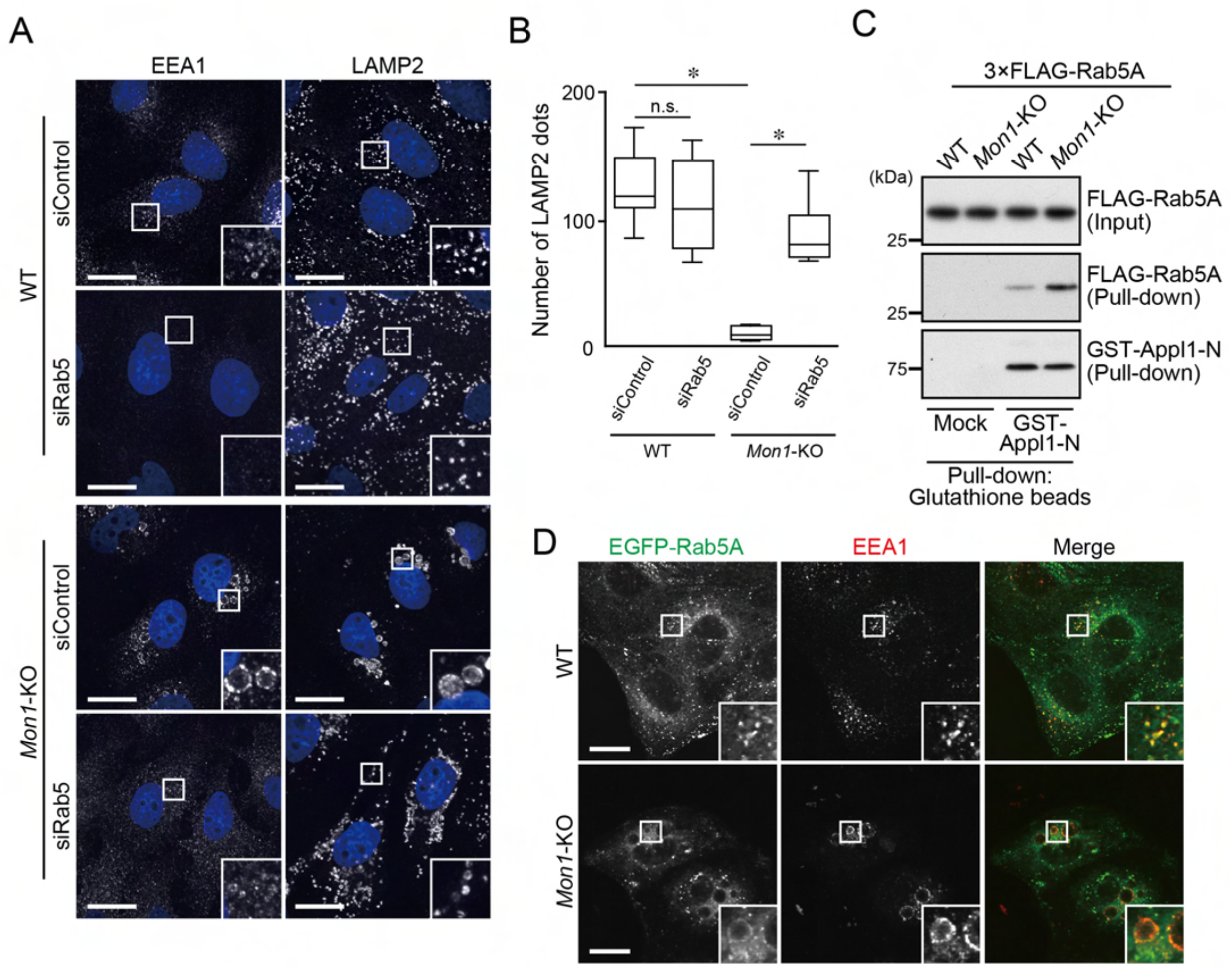
Formation of enlarged endosomes in the *Mon1*-KO is Rab5-dependent. (A) WT and *Mon1*-KO cells were transfected with control siRNA (siControl) or *Rab5A/B/C* siRNA (siRab5). Three days after transfection, the cells were immunostained with anti-LAMP2 antibody. Insets show magnified views of the boxed areas. Scale bars, 20 μm. (B) The numbers of LAMP2-positive dots per cell (n = 15 cells) were counted. The solid bars, boxes, and whiskers indicate the median, interquartile range (25th to 75th percentile), and upper and lower quartiles, respectively, of the values. The statistical analyses were performed by one-way ANOVA and Tukey’s test. *, *P* <0.001. (C) The amount of active, GTP-bound Rab5 in the WT and *Mon1*-KO cells was analyzed by an effector-based pull-down assay with beads coupled to GST-Appl1-N (Zhu et al., 2007). (D) Immunostaining of EEA1 in WT and *Mon1*-KO cells stably expressing EGFP-Rab5A. Insets show magnified views of the boxed areas. Scale bars, 20 μm.

### Comprehensive screening for a candidate(s) Rab5-GAP that regulates endosome maturation

To identify a candidate Rab5-GAP(s) that regulates endosome maturation, we first focused on the protein family members that share a TBC (Tre-2/Bub2/Cdc16)-domain, most of which are known to function as Rab-GAP domains (Fukuda, 2011; Frasa et al., 2012). More than 40 TBC-domain-containing proteins (simply referred to as TBC proteins below) are present in mammals, and although some of them have been shown to exhibit GAP activity toward Rab5 *in vitro* (Xiao et al., 1997; Pei et al., 2002; Chamberlain et al., 2004; Haas et al., 2005; Li et al., 2009; Lachmann et al., 2012; Rao et al., 2021; Yan et al., 2021), no attempt has ever been made to comprehensively screen for Rab5-GAPs in living cells or *in vivo*. If hyperactivation of Rab5 in *Mon1*-KO cells is the primary cause of the greatly enlarged endosomes/lysosomes, forced inactivation of Rab5 by overexpressing TBC/Rab5-GAP proteins should decrease the size of the endosomes/lysosomes, the same as *Rab5* knockdown did (Fig. 2A and 2B). We therefore performed a comprehensive screening for Rab5-GAPs by overexpressing each of the 42 TBC proteins with an EGFP-tag in *Mon1*-KO cells (Fig. S3A). The results showed that overexpression of only two TBC proteins, a known Rab5-GAP, RUTBC3 (Haas et al., 2005; also known as SGSM3), and a previously less characterized TBC protein, TBC1D18, were able to rescue the *Mon1*-KO phenotypes: the enlarged lysosomes in *Mon1*-KO cells became smaller and the number of LAMP2-positive dots increased significantly (Fig. 3A, 3B, and 3C). These rescue effects are most likely to have been caused by inactivation of Rab5 via the TBC domains, because overexpression of the GAP-activity-deficient RUTBC3(RA) and TBC1D18(RK) mutants did not rescue the phenotypes of *Mon1*-KO cells (Fig. 3A and 3C).

**Figure 3.**
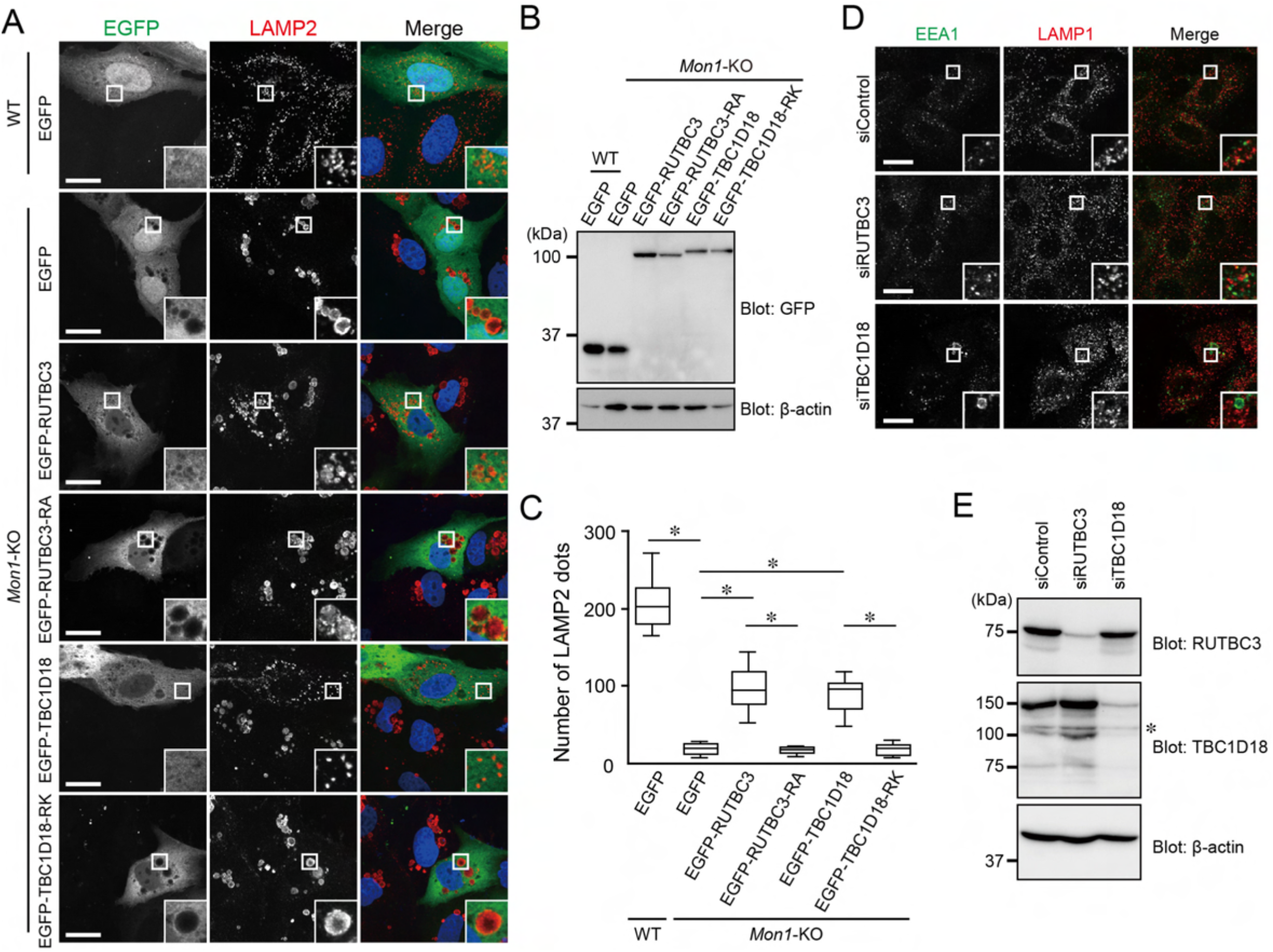
Identification of TBC1D18 as a candidate GAP for Rab5. (A) Immunostaining of LAMP2 in WT cells transiently expressing EGFP or *Mon1*-KO cells transiently expressing EGFP or the EGFP-tagged TBC proteins indicated (see also Fig. S3A). Insets show magnified views of the boxed areas. Scale bars, 20 μm. (B) Lysates of the cells shown in (A) were analyzed by immunoblotting with the antibodies indicated. (C) The numbers of LAMP2-positive dots per cell (n = 15 cells) were counted. The solid bars, boxes, and whiskers indicate the median, interquartile range (25th to 75th percentile), and upper and lower quartiles, respectively, of the values. The statistical analyses were performed by one-way ANOVA and Tukey’s test. *, *P* <0.001. (D) WT cells were transfected with control siRNA (siControl), *RUTBC3* siRNA (siRUTBC3), or *TBC1D18* siRNA (siTBC1D18). Three days after transfection, the cells were immunostained with anti-LAMP1 and anti-EEA1 antibodies. Insets show magnified views of the boxed areas. Scale bars, 20 μm. (E) Lysates of the cells shown in (D) were analyzed by immunoblotting with the antibodies indicated. The asterisk indicates non-specific bands.

Next, we investigated whether endogenous RUTBC3 and TBC1D18 are actually involved in the formation and maturation of endosomes/lysosomes in WT MDCK cells by transfecting them with *RUTBC3* or *TBC1D18* siRNAs and immunostaining them for EEA1 and LAMP1. Depletion of RUTBC3 did not alter either the distribution or size of endosomes/lysosomes (Fig. 3D and 3E), and the *RUTBC3*-KO cells appeared to contain normal endosomes/lysosomes (Fig. S3B and S3C). In sharp contrast, swollen early endosomes and intact lysosomes were observed in the TBC1D18-depleted cells (Figs. 3D, 3E and S3D), which partially resembled the *Mon1*-KO phenotypes (Fig. S1B). We also tried to generate *TBC1D18*-KO cells, but did not succeed: at least one allele of *TBC1D18* was always retained in the sgRNA-expressing cells (data not shown), the same as we had observed in *Rab5*-KO cells in a previous study (Homma et al., 2019), suggesting that TBC1D18 is essential for cell survival and/or growth, the same as Rab5 is. We therefore selected TBC1D18 as the prime candidate for the Rab5-GAP in endosome maturation.

### TBC1D18 functions as a Rab5-GAP both *in vitro* and in cultured cells

Because nothing was known about the function of TBC1D18 (also known as RabGAP1L) in terms of Rab5 regulation, we performed *in vitro* GAP assays with purified components to demonstrate inactivation of Rab5 by TBC1D18. As shown in Fig. 4A and 4B, the amount of GTP bound to Rab5A, which was monitored by luminescence, was found to significantly decrease in a time-dependent manner in the presence of TBC1D18 when compared to the control BSA. Importantly, TBC1D18 did not display any GAP activity toward Rab4A or Rab7 under our experimental conditions. In contrast to these findings in the present study, it has previously been reported that TBC1D18 acts as a GAP for Rab22A, another early endosomal Rab (Itoh et al., 2006; Qu et al., 2016), implying that the phenotypes observed in *Mon1*-KO cells are attributable to Rab22A inactivation by TBC1D18 rather than Rab5 inactivation. To investigate this implication, we transfected *Mon1*-KO cells with *Rab22A* siRNAs and examined them for LAMP2 immunofluorescence. However, the results showed that, unlike Rab5A depletion, Rab22A depletion did not rescue the enlarged lysosome phenotypes in *Mon1*-KO cells, (Fig. S4A and S4B). Moreover, no GAP activity of TBC1D18 toward Rab22A was not detected under our experimental conditions (Fig. S4C). The apparent discrepancy may have been attributable to the difference in substrates used: recombinant untagged Rab22A in this study as opposed to GST-Rab22A in the previous study (Itoh et al., 2006). Actually, N-terminal tagging of EGFP to certain Rabs is now known to distort their functions (Homma and Fukuda, 2016; Oguchi et al., 2020), suggesting that using GST-Rab5A to perform GAP assays is inappropriate.

**Figure 4.**
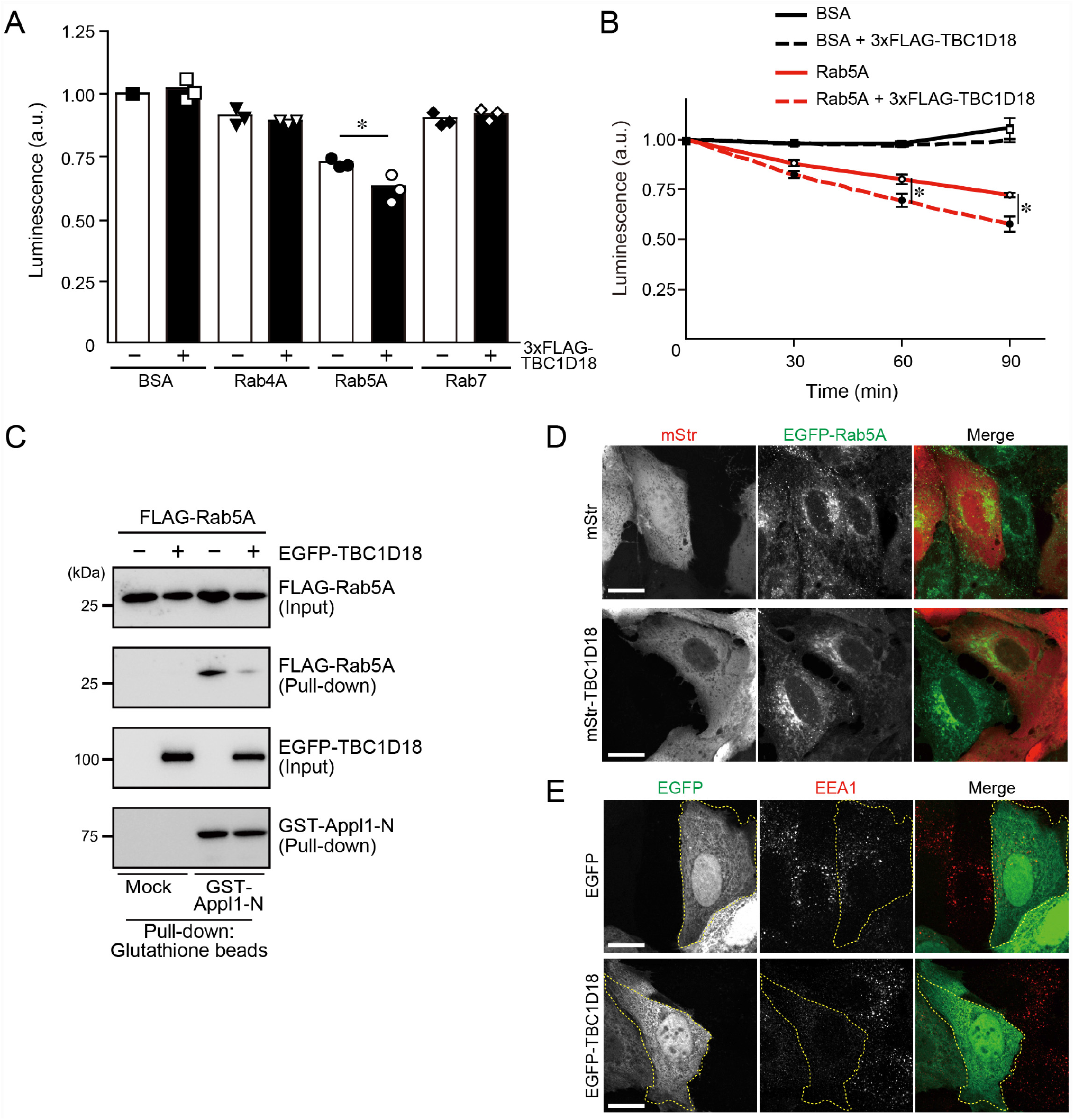
TBC1D18 acts as a Rab5-GAP both *in vitro* and in cultured cells. (A) Quantification of the luminescence, which represents the amount of GTP remaining after GTP hydrolysis by BSA (negative control), Rab4A, Rab5A, and Rab7 in the presence or absence of 3×FLAG-TBC1D18 for 90 min. The statistical analyses were performed by one-way ANOVA and Tukey’s test. *, *P* <0.05. Data are expressed as means and SEM of the data obtained in three independent experiments. (B) Quantification of the luminescence, which represents the amount of GTP remaining after GTP hydrolysis by BSA and Rab5A in the presence or absence of 3×FLAG-TBC1D18 for the times indicated. The statistical analyses were performed by two-way ANOVA and Tukey’s test. *, *P* <0.01. Data are expressed as means and SEM of the data obtained in three independent experiments. (C) The amount of active, GTP-bound Rab5 in WT cells stably expressing EGFP or EGFP-TBC1D18 was analyzed by an effector-based pull-down assay with beads coupled to GST-Appl1-N. (D) Typical images of WT cells stably expressing EGFP-Rab5A and transiently expressing mStr or mStr-TBC1D18. Scale bars, 20 μm. (E) Immunostaining of EEA1 in WT cells transiently expressing EGFP or EGFP-TBC1D18. The cells expressing the protein indicated are outlined with dotted lines.

To further assess whether TBC1D18 has the ability to inactivate Rab5 (i.e., Rab5-GAP activity) even in cultured cells, we performed active Rab5 pull-down assays with GST-Appl1-N in WT cells stably expressing EGFP-TBC1D18, the same as shown in Fig. 2C. The results showed that the amount of active Rab5 in the EGFP-TBC1D18-expressing cells was clearly reduced in comparison with the control EGFP-expressing cells (Fig. 4C), indicating that TBC1D18 actually acts as a Rab5-GAP in cultured cells. Consistent with this finding, the dotted signals of Rab5A and EEA1, a known Rab5 effector, had almost completely disappeared in the WT cells expressing TBC1D18 (Fig. 4D and 4E), suggesting suppression of Rab5 activation by TBC1D18.

### TBC1D18 is involved in endosome maturation through association with Mon1

Since Rab5 is required for endosome fusion and/or maturation (Langemeyer et al., 2018), we further evaluated the impact of TBC1D18 depletion on endosome maturation by performing 488-EGF uptake and EGFR degradation assays. After incubating cells with 488-EGF for 20 min, we immunostained them for EEA1 and LAMP2. The results showed that the EGF-positive dots in the TBC1D18-depleted cells were not co-localized with EEA1 but that the colocalization between EGF-positive dots between LAMP2 was unaffected (Fig. 5A and 5B). Moreover, the same as in Mon1-depleted cells (Fig. 1D), the TBC1D18-depleted cells exhibited attenuated EGFR degradation (Fig. 5C), indicating that TBC1D18 is involved in endosome maturation.

**Figure 5.**
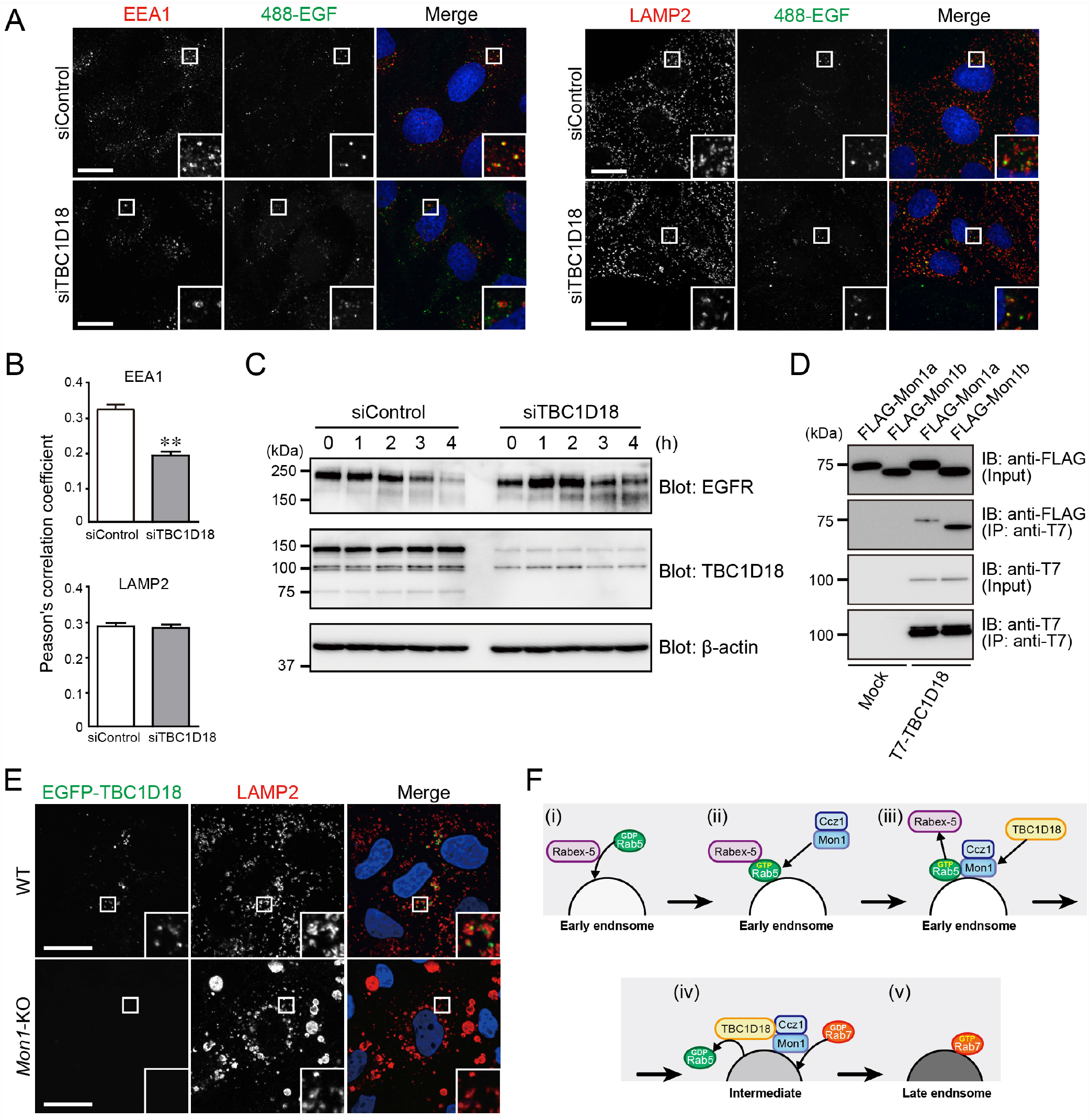
TBC1D18 interacts with Mon1a/b. (A) WT cells that had been treated with the siRNAs indicated were incubated with 488-EGF for 20 min, and then immunostained for EEA1 and LAMP2. Insets show magnified views of the boxed areas. Scale bars, 20 μm. (B) Quantification of Pearson’s correlation coefficients in (A). The data obtained in the cells (n = 15) were analyzed by Student’s *t*-test. **, *P* <0.001. (C) Degradation of EGFR after exposure to 200 ng/mL EGF in WT cells that had been treated with the siRNAs indicated. Their cell lysates were analyzed by immunoblotting with the antibodies indicated. (D) T7-tagged TBC1D18 and FLAG-tagged Mon1a/b were co-expressed in COS-7 cells, and their associations were analyzed by co-immunoprecipitation assays with anti-T7 tag antibody-conjugated agarose beads followed by immunoblotting with the antibodies indicated. (E) After permeabilization with digitonin, WT and *Mon1*-KO cells transiently expressing EGFP-TBC1D18 were fixed and immunostained with anti-LAMP2 antibody. Insets show magnified views of the boxed areas. Scale bars, 20 μm. (F) A new model of Rab5- and Rab7-mediated endosome maturation (see Discussion for details).

Finally, we attempted to determine the functional relationship between Mon1 and TBC1D18 during endosome maturation. One of the simplest relationships would be an association between these two molecules, and the results of co-immunoprecipitation assays indicated that TBC1D18 actually interacts with both Mon1a and Mon1b (Fig. 5D). We then investigated the localization of TBC1D18 in WT and *Mon1*-KO cells. Since TBC1D18 mostly exhibited cytosolic localization (Fig. 3A), which precluded a specific organelle association, we permeabilized the cells with digitonin prior to fixation to remove cytosolic TBC1D18. As shown in Fig. 5E, dotted signals of EGFP-TBC1D18 were observed in close proximity to the LAMP2-positive dots in the WT cells, whereas virtually no EGFP-TBC1D18 signals were observed in the *Mon1*-KO cells, indicating that Mon1 is required for the organelle localization of TBC1D18.

## DISCUSSION

In the present study, we discovered a novel Rab5-GAP, TBC1D18, that mediates endosome maturation in a Mon1-depenedent manner. Since TBC1D18 interacted with Mon1, a Rab7-GEF component, and its organelle localization (i.e., close proximity to LAMP2-positive compartments) was Mon1-dependedent (Fig. 5D and 5E), we propose the following Mon1–TBC1D18-mediated endosome maturation model in mammalian cells (Fig. 5F): (i) Rabex-5, a Rab5-GEF, activates Rab5 on early endosomes. (ii) Rab5 then recruits the Mon1–Ccz1 complex to early endosomes as a Rab5 effector (Kinchen & Ravichandran, 2011). (iii) The recruited complex in turn “recruits TBC1D18, which inactivates Rab5” (this study) and promotes dissociation of Rabex-5 from early endosomes (Poteryaev et al., 2010). (iv) The recruited TBC1D18 then inactivates Rab5, and the Mon1–Ccz1 complex activates Rab7 as a Rab7-GEF, both of which promote efficient conversion from early endosomes to late endosomes (v). Thus, Mon1 is likely to function as a hub for endosome maturation that coordinates the Rab5-GEF activity of Rabex-5, the Rab5-GAP activity of TBC1D18, and its own Rab7-GEF activity.

Although Mon1 plays dual roles in Rab5 inactivation by promoting Rabex-5 dissociation and TBC1D18 recruitment and Rab7 activation via its GEF activity, the Rab5 inactivation appears to be more important than the Rab7 activation in terms of endosome maturation. Actually, *Mon1*-KO cells showed severe defects in endosome maturation (i.e., enlarged lysosomes and attenuated EGFR degradation), whereas *Rab7*-KO cells showed rather mild phenotypes (Fig. 1A and 1D). Moreover, the phenotypes of the *Mon1*-KO cells resembled those of Rab5A-QL-expressing cells (Fig. S2A). Since Mon1 is recruited to early endosomes via Rab5 even in *Rab7*-KO cells, Rab5 must be inactivated by the Mon1–TBC1D18 axis, which would maintain endo-lysosomal morphology and function to some extent. Hyperactivation of Rab5 is known to facilitate homotypic fusion between early endosomes (Stenmark et al., 1994), and in the presence of Rab5 these enlarged early endosomes may not mature into late endosomes. Nevertheless, the enlarged EEA1-positive structures in *Mon1*-KO cells and Rab5-QL-expressing cells were also positive for LAMP2, suggesting that enlarged early endosomes can fuse with lysosomes before fully maturing into late endosomes. Although these hybrid structures were LysoTracker- and Magic Red-positive, lysosomal degradation activity as monitored by EGFR degradation and DQ-BSA was clearly inhibited (Figs 5C and S1E). Thus, proper inactivation of Rab5 before fusion with lysosomes is necessary to achieve efficient degradation of endocytic cargos.

In addition to TBC1D18, another TBC protein, RUTBC3, has previously been shown to function as a Rab5-GAP (Haas et al., 2005). The results of our initial comprehensive screening in *Mon1*-KO cells also pointed to RUTBC3 as a possible Rab5-GAP (Fig. 3A). Unlike TBC1D18, however, *RUTBC3* KO or knockdown did not affect endosome formation (Fig. S3C and S3D). Although human and mouse *TBC1D18* and *RUTBC3* mRNAs are ubiquitously expressed in a variety of tissues (NCBI data base; gene IDs: 9910 and 27352, respectively), it is still possible that Rab5 uses different Rab5-GAPs in different cell-types and/or different tissues. Alternatively, the existence of multiple Rab5-GAPs, including TBC1D18 and RUTBC3 (Haas et al., 2005; Li et al., 2009; Rao et al., 2021; Yan et al., 2021), may ensure the regulation of diverse endocytic pathways such as receptor-mediated endocytosis, fluid-phase endocytosis (pinocytosis), and phagocytosis, in individual cells.

The transition from Rab5 to Rab7 during endosome maturation is a mechanism that is common to all eukaryotic cells, and the Mon1–Ccz1 complex has been retained during evolution (Borchers et al., 2021). TBC1D18, however, is conserved in vertebrates alone, implying that the coordinated regulation of endosome maturation by TBC1D18 and Mon1 may have been evolutionarily acquired in vertebrates. Nevertheless, since TBC proteins are widely present in invertebrates (Fukuda, 2011), including in fungi and plants, it is still possible that Mon1-dependent Rab5 inactivation is the common mechanism and that other TBC proteins function during endosome maturation in invertebrates (i.e., convergent evolution in terms of Rab5 inactivation). Further research will be necessary to elucidate whether Mon1-dependent Rab5 inactivation is important for endosome maturation in other invertebrate species.

In summary, we succeeded in identifying a novel Rab5-GAP, TBC1D18, that mediates endosome maturation, and we have proposed a new model in which Mon1 recruits TBC1D18 to endosomes/lysosomes, which accelerates a transition from Rab5 to Rab7 during endosome maturation by inactivating Rab5 and activating Rab7. Intriguingly, Mon1/TBC1D18-mediated Rab5 inactivation is more important than Mon1-mediated Rab7 activation during endosome maturation, because the defects in endosome maturation were more severe after Rab5 hyperactivation than after Rab7 inactivation. Since both Mon1 and TBC1D18 appear to localize mostly in the cytosol, the spatiotemporal regulation of their activities and the precise mechanism of their targeting to Rab5-localzied endosomes during endosome maturation are important issues that need to be addressed next in the future studies.

## MATERIALS AND METHODS

### Materials

Rabbit polyclonal antibodies against Rab5B/C, Rab7, and Rab22A were prepared as described previously (Mrozowska & Fukuda, 2016). The following primary antibodies were obtained commercially: anti-Mon1a goat polyclonal antibody (Biorbyt, San Francisco, CA; #orb20562), anti-Mon1b mouse polyclonal antibody (Abnova, Taipei Taiwan; #H00022879-B01P), anti-CD107b (LAMP2) mouse monoclonal antibody (Thermo Fisher Scientific, Waltham, MA; #MA5-28269), anti-LAMP1 rabbit polyclonal antibody (Abcam, Cambridge, UK; #ab24170), anti-Rab5 rabbit polyclonal antibody (Thermo Fisher Scientific; #PA5-29022), anti-EGFR sheep polyclonal antibody (Fitzgerald, Acton, MA; #20-ES04), anti-ß-actin mouse monoclonal antibody (Applied Biological Materials, Richmond, BC, Canada; #G043), anti-EEA1 mouse monoclonal antibody (BD Biosciences, San Jose, CA; #610456), anti-GM130 mouse monoclonal antibody (BD Biosciences; #610823), anti-LBPA mouse monoclonal antibody (Echelon, Santa Clara, CA; #Z-SLBPA), anti-SGSM3 (RUTBC3) rabbit polyclonal antibody (Proteintech, Rosemont, IL; #20825-1-AP), anti-RABGAP1L (TBC1D18) rabbit polyclonal antibody (Proteintech; #13894-1-AP), horseradish peroxidase (HRP)-conjugated anti-GFP polyclonal antibody (MBL, Nagoya, Japan; #598-7), HRP-conjugated anti-FLAG mouse monoclonal antibody (Sigma-Aldrich, St. Louis, MO; #A8592-1MG), HRP-conjugated anti-T7 mouse monoclonal antibody (Merck Biosciences Novagen, Darmstadt, Germany; #69048), and HRP-conjugated anti-GST (Z-5) rabbit polyclonal antibody (Santa Cruz Biotechnology, Dallas, TX; #sc-459). HRP-conjugated anti-mouse IgG goat polyclonal antibody (SouthernBiotech, Birmingham, AL; #1031-05), HRP-conjugated anti-rabbit IgG donkey polyclonal antibody (GE Healthcare, Chicago, IL; #NA934V), HRP-conjugated anti-sheep IgG donkey polyclonal antibody (Chemicon International, Temecula, CA; #AP184P), HRP-conjugated anti-goat IgG rabbit polyclonal antibody (Chemicon International; #AP106P), Alexa Fluor 488^+^-conjugated anti-mouse IgG goat polyclonal antibody, Alexa Fluor 555^+^-conjugated antirabbit IgG goat polyclonal antibody, and Alexa Fluor 555^+^-conjugated anti-mouse IgG goat polyclonal antibody (Thermo Fisher Scientific; #32723, #32732, and #A32727, respectively) were also obtained commercially.

Magic Red™ Cathepsin B Assay (#937) was purchased from Immunochemistry Technologies, Bloomington, MN. LysoTracker™ Red DND-99 (#L7528), DQ™ Red BSA (#D12051), and Alexa Fluor 488 EGF complex (488-EGF) (#E13345) were from Thermo Fisher Scientific.

### cDNA cloning and plasmid constructions

cDNA encoding mouse Mon1a was prepared as described previously (Yasuda et al., 2016), and the mouse Mon1b cDNA was amplified from the Marathon-Ready adult mouse brain and testis cDNAs (Clontech/Takara Bio, Shiga, Japan) by conventional PCR techniques. The Mon1a and Mon1b cDNAs obtained were subcloned into the pMRX-puro (Saitoh et al., 2003) and pEF-FLAG vectors (Fukuda et al., 1999). cDNAs encoding mouse Rab4A Rab5A, Rab5A-Q79L, Rab7, Rab7-Q67L, and Rab22A were prepared as previously described (Itoh et al., 2006). The Rab4A, Rab5A, Rab7, and Rab22A cDNAs were subcloned into the pGEX-4T-3-gl vector, which was constructed by using specific oligonucleotides in this study for recombinant protein purification. This vector contains a Gly linker sequence (PGIS<u>GGGGG</u>S) just downstream of its thrombin recognition sequence for efficient thrombin digestion. The Rab5A, Rab5A-Q79L (QL), and Rab7-Q67L (QL) cDNAs were also subcloned into the pMRX-puro-EGFP vector. cDNA encoding the N-terminal region (5–419 amino acids) of mouse Appl1 was amplified from the Marathon-Ready adult mouse brain and testis cDNAs by PCR and subcloned into the pGEX-4T-3 vector. pEGFP-C1-TBC vectors were prepared as described previously (Ishibashi et al., 2009). RUTBC3-R165A (RA) and TBC1D18-R588K (RK) point mutants were prepared with specific oligonucleotides by the standard molecular biology techniques. TBC1D18-WT and -RK cDNAs were subcloned into the pMRX-puro-EGFP vector, pMRX-puro-3×FLAG vector, which was constructed by using specific oligonucleotides in this study, pmStr-C1 vector (Ohbayashi et al., 2012), and/or pEF-T7 vector (Fukuda et al., 1999). pEF-T7-RILP was prepared as described previously (Matsui et al., 2012).

### Cell cultures and transfection

COS-7 cells and MDCK cells (WT and Rab7-KO MDCK-II cells; RIKEN BioResource Research Center, Cat# RCB5107 and RCB5148, respectively) (Homma et al., 2019) were cultured at 37°C in Dulbecco’s modified Eagle’s medium (DMEM) (Fujifilm Wako Pure Chemical, Osaka, Japan) supplemented with 10% fetal bovine serum, 100 units/mL penicillin G, and 100 μg/mL streptomycin in a 5% CO2 incubator. One day after plating, cells were transfected with plasmids and siRNA oligonucleotides by using Lipofectamine 2000 (Thermo Fisher Scientific) and Lipofectamine RNAiMAX (Thermo Fisher Scientific), respectively, each according to the manufacturer’s instructions.

### Generation of KO cells

*Mon1*-KO and *RUTBC3*-KO MDCK cells were generated by using the CRISPR/Cas9 system as described previously (Homma et al., 2019). The single guide RNA sequences targeting dog Mon1a, Mon1b, and RUTBC3 were 5’-ACAAGGTAGTATTCGTGCGC-3’, 5’-ACTCTGCACGAAGGACACGA-3’, and 5’-AAGCCTGCTAGAAGTGGGGT-3’, respectively.

### Retrovirus production and infection into MDCK cells

Retroviruses were produced in Plat-E cells as described previously (Homma et al., 2019). The virus-containing medium was added to the culture medium of MDCK cells in the presence of 8 μg/mL polybrene, and after 24 h the transformants were selected with 2 μg/mL puromycin (Merck) for 24–48h.

### Recombinant protein purification

GST-Appl1-N, GST-gl-Rab4A, GST-gl-Rab5A, GST-gl-Rab7, GST-gl-Rab22A, and control GST alone were expressed in *E. coli* JM109 and purified with glutathione-Sepharose beads (GE Healthcare) by the standard protocol. To assay GAPs for purified Rabs, GST-tag of GST-Rabs were cleaved by thrombin digestion, and the cleaved GST and thrombin were removed with glutathione-magnetic beads (GenScript, Piscataway, NJ) and benzamidine-Sepharose beads (GE Healthcare), respectively.

3×FLAG-TBC1D18 was transiently expressed in COS7 cells, and the cells were lysed with lysis buffer #1 (50 mM HEPES-KOH, pH7.2, 150 mM NaCl, 1% Triton X-100, and 1 mM EDTA) containing a phosphatase inhibitor cocktail (Nacalai Tesque, Kyoto, Japan) and protease inhibitor cocktail (Roche, Basel, Switzerland). After centrifugation at 20,380×*g* for 10 min, the supernatants obtained were incubated with gentle rotation for 1 h at 4°C with anti-FLAG tag-antibody-conjugated magnetic beads (Sigma-Aldrich). The beads were then washed three times with lysis buffer #1 and incubated with gentle rotation for 30 min at 4°C with 50 μL of an elution buffer (50 mM HEPES-KOH, pH7.2, 150 mM NaCl, 500 μg/mL 3×FLAG peptide [Sigma-Aldrich]). The final supernatant containing 3×FLAG-TBC1D18 was transferred to a fresh tube.

### Immunoblotting

Protein extracts obtained from cells that had been lysed with an SDS sample buffer were boiled for 10 min. The protein samples were separated by SDS-PAGE and transferred to PVDF membranes (Merck Millipore, Burlington, MA) by electroblotting. The membranes were blocked for 30 min at room temperature with 1% skimmed milk in PBS containing 0.1% Tween-20 (PBS-T) and incubated overnight at 4°C with a primary antibody diluted in the blocking buffer. After washing three times with PBS-T, they were incubated for 90 min at room temperature with an appropriate HRP-conjugated secondary antibody. Chemiluminescence signals were visualized with the Immobilon Western Chemiluminescent HRP substrate (EMD Millipore, Burlington, MA) and detected with a chemiluminescence imager (ChemiDoc Touch; Bio-Rad, Hercules, CA).

### Immunofluorescence analysis

Cells grown on coverslips were fixed with 4% paraformaldehyde for 10 min then washed them with PBS three times. The fixed cells were permeabilized for 5 min at room temperature with 50 μg/mL digitonin (Sigma-Aldrich) in PBS and then blocked for 30 min at room temperature with 3% BSA in PBS. To remove cytosolic components of EGFP-TBC1D18 in Fig. 5E, the cells were permeabilized with digitonin was performed before fixation. The permeabilized cells were incubated for 1 h with a primary antibody, washed with PBS, and then incubated for 1 h with Alexa-labeled anti-mouse or rabbit IgG secondary antibody. After washing the cells with PBS, the coverslips were mounted on glass slides with Prolong Diamond (Thermo Fisher Scientific, #P36961) and with DAPI. All procedures were carried out at room temperature. Confocal fluorescence images were obtained through a confocal fluorescence microscope (Fluoview 1000; Olympus, Tokyo, Japan) equipped with a Plan-Apochromat 100×/1.45 oil-immersion objective lens and an electron-multiplying charge-coupled device camera (C9100; Hamamatsu Photonics, Shizuoka, Japan).

### Electron microscopy

Cells were cultured on cell-tight C-2 cell disks (Sumitomo Bakelite, Tokyo, Japan; #MS-0113K) and fixed for 2 h on ice in 2.5% glutaraldehyde (Electron Microscopy Science, Hatfield, PA; #111-30-8) in 0.1 M phosphate buffer, pH 7.4. The cells were washed with 0.1 M phosphate buffer, pH 7.4 three times, postfixed in 1% osmium tetroxide in 0.1 M phosphate buffer, pH 7.4 for 2 h, dehydrated, and embedded in Epon 812 according to the standard procedure. The samples were examined with an H-7100 electron microscope (Hitachi High-Tech Corp., Tokyo, Japan).

### EGFR degradation assay

Cells grown on a 24-well plate were starved for 24 h in serum-free DMEM. After pre-incubation at 37°C for 30 min in the medium containing 100 μg/mL cycloheximide, the cells were exposed to 200 ng/mL EGF for the times indicated in Figs. 1D and 5C. The total EGFR protein level after the EGF treatment was evaluated by immunoblotting.

### GTP-Rab pull-down assays performed with Rab effector domains

For the GTP-Rab7 pull-down assays, T7-tagged RILP, a Rab7 effector, was expressed in COS-7 cells and immobilized to anti-T7 tag-antibody-conjugated agarose beads (EMD Millipore) (Matsui et al., 2012). MDCK-WT and *Mon1*-KO cells were lysed with lysis buffer #1 containing a protease inhibitor cocktail, and the supernatants were incubated with rotation for 1 h at 4°C with beads coupled to T7-RILP. The beads were washed three times with the lysis buffer #1, boiled for 5 min with an SDS sample buffer, and then subjected to SDS-PAGE and immunoblotting analyses.

For the GTP-Rab5 pull-down assays, MDCK cells were lysed with lysis buffer #2 (50 mM HEPES-KOH, pH7.2, 150 mM NaCl, 0.1% NP-40, and 10 mM MgCl2) containing a phosphatase inhibitor cocktail (Nacalai Tesque) and protease inhibitor cocktail (Roche). After centrifugation at 20,380×*g* for 10 min, the supernatants obtained were incubated with rotation for 1 h at 4°C with beads coupled to GST or GST-Appl1-N. The beads were then washed three times with a buffer (50 mM HEPES-KOH, pH 7.2, 150 mM NaCl, and 10 mM MgCl2), boiled for 5 min with an SDS sample buffer, and then subjected to SDS-PAGE and immunoblotting analyses.

### *In vitro* GAP assay

GAP-accelerated GTP hydrolysis of Rab was measured by using a GTPase-Glo™ assay kit (Promega, Madison, WI; #V7681) according to the manufacture’s instruction. In brief, purified recombinant Rab4A, Rab5A, Rab7, or Rab22A (0.45 μg each) with or without 3×FLAG-TBC1D18 (0.033 μg) was suspended in the reaction buffer in the kit. After addition of GTP to a final concentration of 5 μM, the reaction mixture was incubated at room temperature for the times indicated in Fig. 4A and 4B, and in Fig. S4C. The GTPase-Glo™ reagent was added to the reaction mixture to convert the remaining GTP to ATP. The amount of ATP was then detected with luciferase to produce bioluminescence, which was measured with a Victor Nivo Multimode microplate reader (PerkinElmer, Waltham, MA).

### Assay for endocytic cargo uptake

Cells grown on coverslips were exposed to 25 μg/mL DQ^TM^ Red BSA in serum-free DMEM for 6 h or to 2 μg/mL 488-EGF in serum-free DMEM for 20 min. After incubation, the cells were fixed with 4% paraformaldehyde and subjected to immunofluorescence analysis.

### Co-immunoprecipitation assay

COS-7 cells transiently expressing recombinant proteins were lysed for 10 min on ice in lysis buffer #1 containing a phosphatase inhibitor cocktail and protease inhibitor cocktail. After centrifugation at 20,380×*g* for 10 min, the supernatants were incubated with gentle rotation for 1 h at 4°C with anti-T7 tag-antibody-conjugated agarose beads. The beads were washed three times with lysis buffer #1, boiled for 5 min with an SDS sample buffer, and then subjected to SDS-PAGE and immunoblotting analyses.

## Acknowledgments

We thank Kazuyasu Shoji for technical assistance, Dr. Yuta Homma for initial Mon1 experiments, Drs. Toshio Kitamura and Shoji Yamaoka for kindly donating Plat-E cells and pMRX vector, respectively, and all members of the Fukuda laboratory for helpful discussions. This work was supported in part by Grant-in-Aid for Early-Career Scientists 20K15786 (to T. M.) from the Ministry of Education, Culture, Sports, Science and Technology (MEXT) of Japan; Grant-in-Aid for Scientific Research (B) 19H03220 from MEXT (to M. F.); Japan Science and Technology Agency (JST) CREST Grant JPMJCR17H4 (to M. F.); and by Tohoku University Division for interdisciplinary Advanced Research and Education (to S. H.).

## Author contributions

Conceptualization, S. H., T. M., and M. F.; Investigation, S. H., T. M., and Y. S.; Writing – Original Draft, S. H., and M. F.; Writing – Review & Editing, S. H., T. M., Y. S., and M. F.; Funding Acquisition, T. M. and M. F.; Supervision, M. F.

## SUPPLEMENTAL FIGURE LEGENDS

**Figure S1.**
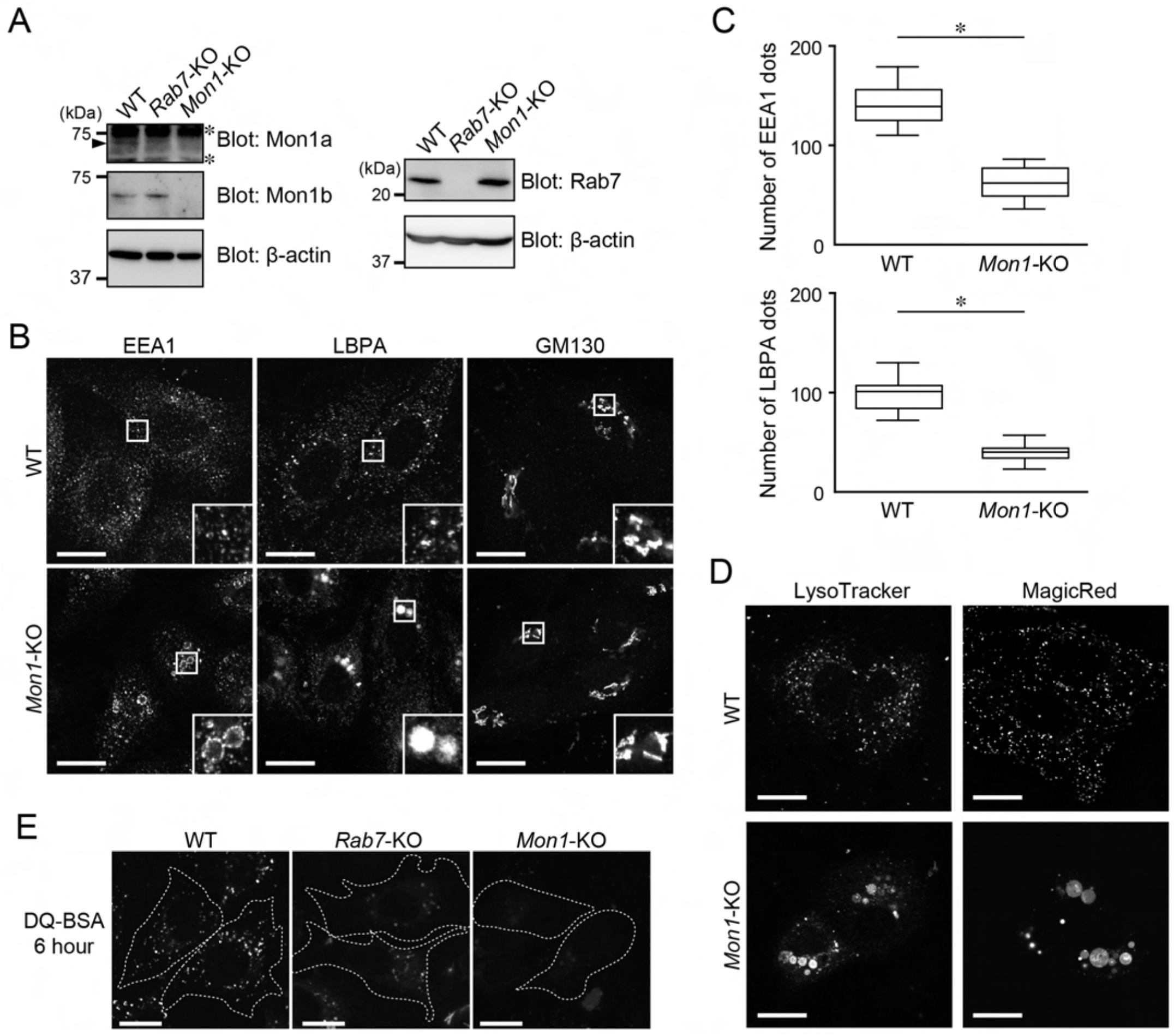
Additional endosome and lysosome phenotypes in *Mon1*-KO cells (related to Figure 1). (A) Lysates of WT, *Rab7*-KO, and *Mon1*-KO cells were analyzed by immunoblotting with the antibodies indicated. The asterisks and arrowhead indicate non-specific bands and Mon1a, respectively. (B) Immunostaining of EEA1, LBPA, and GM130 in WT and *Mon1*-KO cells. Insets show magnified views of the boxed areas. Scale bars, 20 μm. (C) The numbers of EEA1- or LBPA-positive dots per cell (n = 15 cells) were counted. The solid bars, boxes, and whiskers indicate the median, interquartile range (25th to 75th percentile), and upper and lower quartiles, respectively, of the values. The statistical analyses were performed by Student’s *t*-test. *, *P*<0.01. (D) Typical images of WT and *Mon1*-KO cells that had been exposed to LysoTracker and MagicRed. Scale bars, 20 μm. (E) Typical images of WT, *Rab7*-KO, and *Mon1*-KO cells (outlined with dotted lines) that had been exposed to DQ-BSA for 6 h. Cells are outlined with dotted lines. Scale bars, 20 μm.

**Figure S2.**
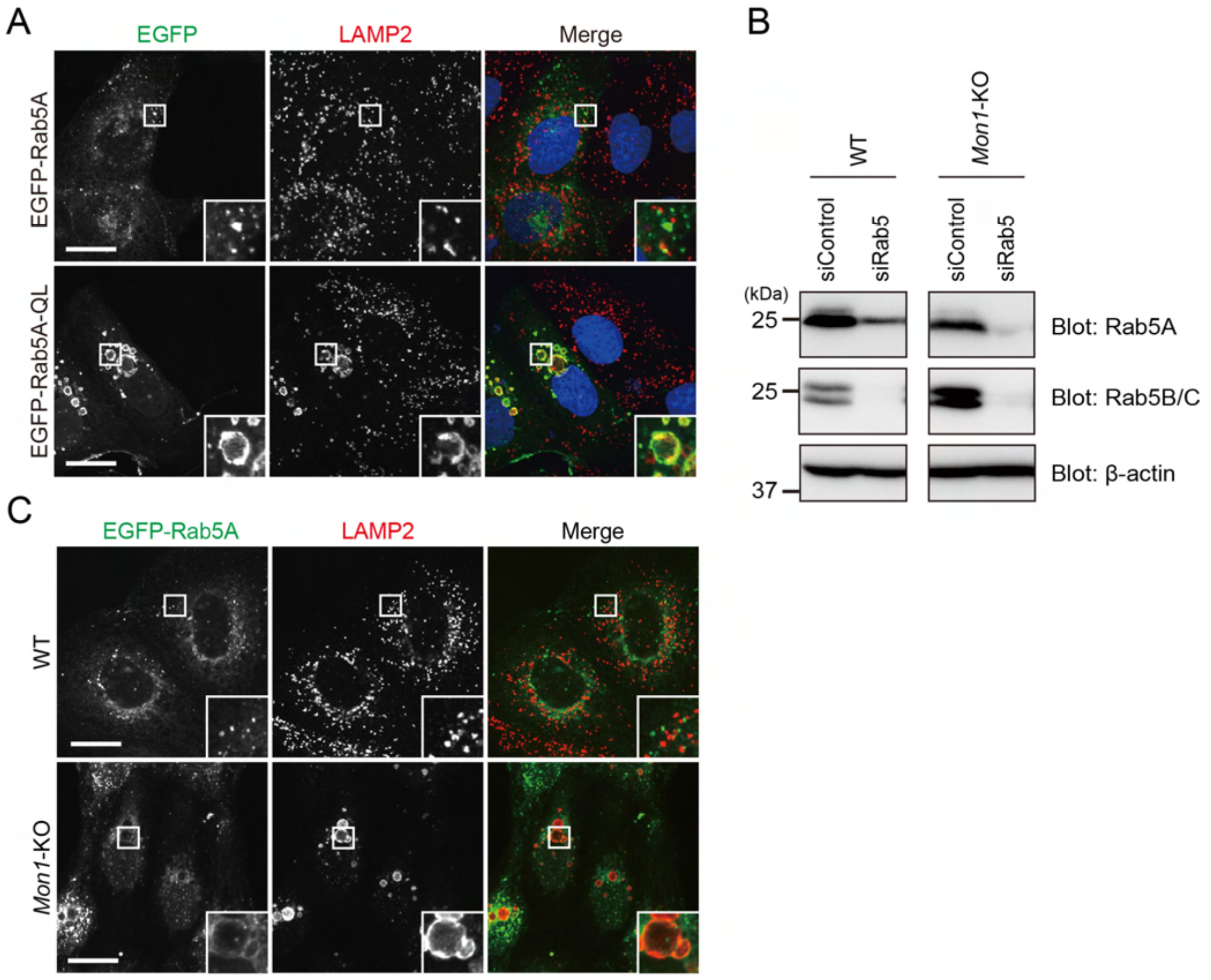
Hyperactivation of Rab5 in *Mon1*-KO cells (related to Figure 2). (A) Immunostaining of LAMP2 in WT cells transiently expressing EGFP-Rab5A or EGFP-Rab5A-QL. Insets show magnified views of the boxed areas. Scale bars, 20 μm. (B) Lysates of the cells shown in Fig. 2A were analyzed by immunoblotting with the antibodies indicated. (C) Immunostaining of LAMP2 in WT and *Mon1*-KO cells transiently expressing EGFP-Rab5A. Insets show magnified views of the boxed areas. Scale bars, 20 μm.

**Figure S3.**
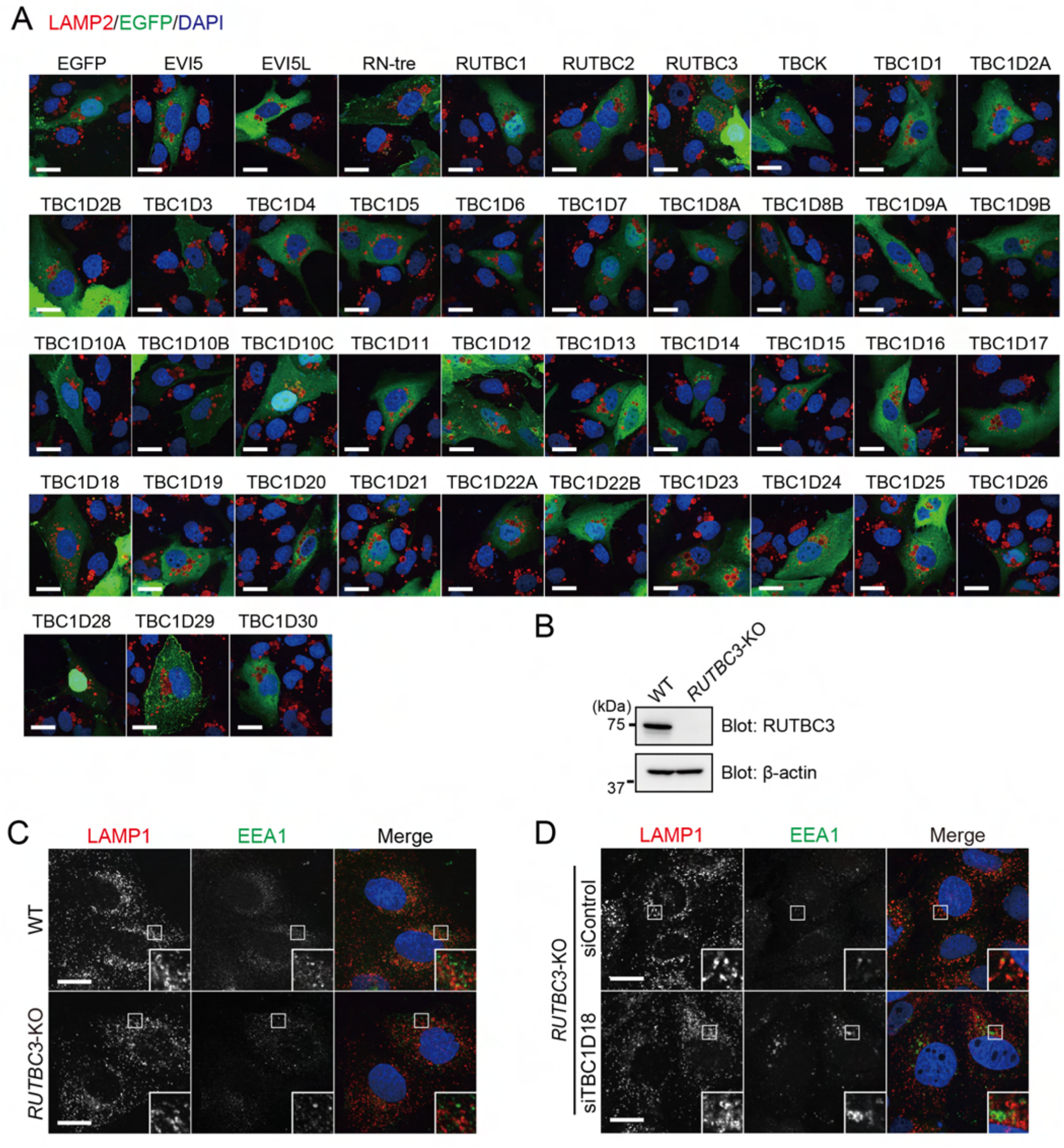
Summary of the comprehensive screening of 42 TBC proteins and analysis of *RUTBC3*-KO phenotypes (related to Figure 3). (A) Immunostaining of LAMP2 in *Mon1*-KO cells transiently expressing EGFP-TBC proteins. Scale bars, 20 μm. (B) Lysates of WT and *RUTBC3*-KO cells were analyzed by immunoblotting with the antibodies indicated. (C) Immunostaining of EEA1 and LAMP1 in WT and *RUTBC3*-KO cells. Insets show magnified views of the boxed areas. Scale bars, 20 μm. (D) *RUTBC3*-KO cells were transfected with control siRNA (siControl) or *TBC1D18* siRNA (siTBC1D18). Three days after transfection, the cells were immunostained with anti-LAMP1 and anti-EEA1 antibodies. Insets show magnified views of the boxed areas. Scale bars, 20 μm.

**Figure S4.**
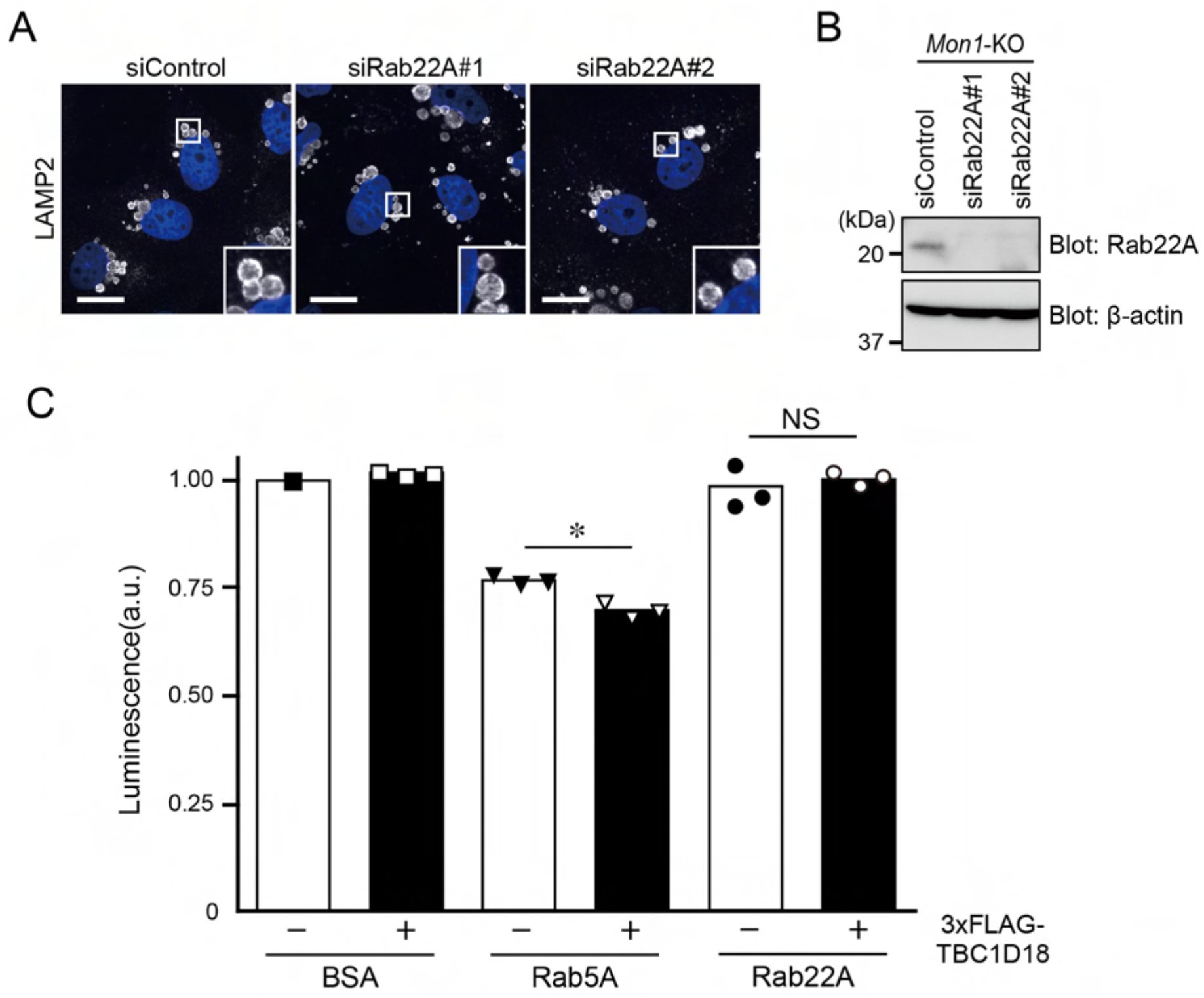
Rab22A is unlikely to be involved in the phenotypes of *Mon1*-KO cells (related to Figure 4). (A) WT cells were transfected with control siRNA (siControl) or *Rab22A* siRNAs (siRab22A#1 and #2). Three days after transfection, the cells were immunostained with anti-LAMP2 antibody. Insets show magnified views of the boxed areas. Scale bars, 20 μm. (B) Lysates of the cells shown in (A) were analyzed by immunoblotting with the antibodies indicated. (C) Quantification of the luminescence, which represents the amount of GTP remaining after GTP hydrolysis by BSA (negative control), Rab5A, and Rab22A for 90 min in the presence or absence of 3×FLAG-TBC1D18. The statistical analyses were performed by one-way ANOVA and Tukey’s test. *, *P* <0.05. Data are expressed as means and SEM of the data obtained in three independent experiments.

